# Tree species composition and nutrient uptake influence soil nutrient availability, even under contrasted levels of water availability

**DOI:** 10.64898/2026.10.29.761915

**Authors:** Tania L. Maxwell, Nicolas Fanin, Alison D. Munson, Catherine Lambrot, Pierre Trichet, Frederic Bernier, Laurent Severin, Laurent Augusto

## Abstract

Temporal asynchrony within a vegetation period, such as when mixing trees with contrasting leaf phenology (i.e. deciduous and evergreen species), may play an important role in accelerating belowground nutrient cycling in mixed-species forests. However, forests are increasingly impacted by drought, and how mixing effects may be modified by changes in water availability is unknown. Using an 11-year-old experimental plantation in southwestern France, we tested the interactive effect of stand composition (monocultures of each *Pinus pinaster* and *Betula pendula*, and the mixture of the two species) and water availability (*via* irrigation) on various belowground processes. We quantified soil nitrogen (N) and phosphorus (P) availability with ion-exchange resins in the growing vs dormant periods, net mineralization with an *in situ* soil incubation method every 4-6 weeks from April to September, and N uptake by trees using two *in situ* ^15^N soil labeling campaigns, one at budburst and one at peak leaf expansion in summer. The methods were implemented at several time points, in order to elucidate potential temporal complementarity in the mixture. We found differences in soil nutrient availability between different species compositions, with pure pine generally having higher mineral N, and pure birch having higher available P under high water availability. These differences were unlikely due to increases in net nutrient mineralization, as these rates were similar among treatments. Instead, differences in tree growth and canopy covering between species compositions, and thus different nutrient uptake needs may be the main drivers for the soil nutrient availability trends. The ^15^N labeling experiment revealed a competitive advantage for pine in the mixture at budburst, with higher N uptake than pine in monocultures, without negatively impacting birch uptake. This advantage was then lost once birch’s leaves were out in the early summer. We conclude that the timing of nutrient needs and uptake of species within a mixture, such as between a broadleaf deciduous and an evergreen conifer, may be a stronger driver than mineralization for soil nutrient availability in a mixture, even under two contrasted levels of water availability.

## I Introduction

While forests provide a multitude of ecosystem services (Gamfeldt et al., 2013; Liang et al., 2016; Jactel et al., 2017), they are increasingly threatened by more intense and frequent drought periods (Schwalm et al., 2012; IPCC, 2014), with important negative consequences on tree productivity and survival (Hartmann et al., 2018). Increasing tree diversity has been a well-explored option to mitigate these negative effects (Lebourgeois et al., 2013; Pretzsch et al., 2013) while improving ecosystem services such as nutrient cycling (Richards et al., 2010). Mixed-species forests may be more stable under climatic fluctuations because of species asynchrony and positive species interactions (Jucker et al., 2014; Morin et al., 2014; Kardol et al., 2018), which can enhance nutrient supply (Richards et al., 2010; Zhang et al., 2026) and productivity relative to pure stands (Cardinale et al., 2007; Zhang et al., 2012; Liang et al., 2016; Baeten et al., 2019). Positive effects of tree diversity on soil nutrient availability are often complementarity among the different species (Loreau and Hector, 2001), which may occur through spatial or temporal niche differentiation (Forrester, 2014). Spatial complementarity can arise from differences or soil resource acquisition in rooting depth or soil resource acquisition (Brassard et al., 2013; Altinalmazis-Kondylis et al., 2020), whereas temporal complementarity may result from differences in the timing of resource acquisition and canopy development. Although temporal stability of productivity in mixed forests has been extensively linked to asynchronous canopy dynamics (Forrester, 2014; Morin et al., 2014; del Río et al., 2017; Martin-Blangy et al., 2023), much less is known about temporal complementarity in belowground nutrient cycling within a growing season. Understanding these mechanisms is particularly important because biodiversity–ecosystem functioning relationships may become stronger under water-limited conditions (Ratcliffe et al., 2017), yet we still know little about how changes in water availability influence diversity effects on belowground nutrient cycling.

Mixing trees with different nutrient requirements and timing of resource acquisition may lead to seasonal complementarity of belowground N cycling (Rothe and Binkley, 2001). In general, net N mineralization is expected to be higher in broadleaved-dominated forests than in conifer-dominated forests (Gosz, 1981; Melillo, 1981; Côté et al., 2000), due to the nutrient-rich litter stimulating belowground microbial activity, although other factors such as soil properties, soil moisture or understory communities can also influence mineralization processes (Brierley et al., 2001; Lejon et al., 2005; Augusto et al., 2015). Mixing broadleaved and needle litter has frequently been reported to increase N mineralization beyond predicted values (Chapman et al., 1988; Williams and Alexander, 1991; McTiernan et al., 1997; Salamanca et al., 1998; Gartner and Cardon, 2004; Fyles and Fyles, 2011), as species with nutrient-rich litter may enhance the decomposition and nutrient release of species producing low-quality litter (Chapman et al., 1988; Wardle et al., 1997; Finzi and Canham, 1998). Temporal complementarity may therefore arise through variable rates of nutrient release in mixed compared to monoculture stands throughout the growing season (McTiernan et al., 1997), due to differences in litter qualities and recalcitrance. This complementarity could contribute to a more even release of inorganic nutrients over the growing season, reducing nutrient losses and increasing nutrient availability for plant uptake (McTiernan et al., 1997; Richards et al., 2010). Another potential mechanism is a difference in the timing of activity and nutrient uptake between deciduous and evergreen species, as activity in the latter can start earlier due to foliage that remains functional year-round (Krishna and Chandra Garkoti, 2022). Changes in nutrient phenology when species grow in mixture compared with monoculture may also lead to differences in the timing of nutrient uptake, potentially reducing resource losses from the ecosystem (Knops et al., 2002). However, the extent to which mixing tree species increases nutrient capture through seasonal complementarity remain unclear (Forrester, 2014). Most studies have relied on indirect indicators such as root biomass and nutrient concentrations (Lang and Polle, 2011), soil nutrient availability (Maxwell et al., 2020a), or aboveground nutrient content (Oelmann et al., 2010; Maxwell et al., 2022b), to infer nutrient capture in mixed forests.

Water availability plays an important role in controlling belowground processes, as it regulates soil microbial activity (Elliott et al., 1981; Brockett et al., 2012), with decreased soil water content that generally reduces microbial activity in aerobic conditions (Kreuzwieser and Gessler, 2010). In forests, limited water supply may reduce litter decomposition rates (Zheng et al., 2017), mineralization (Borken and Matzner, 2009), and nitrification (Chen et al., 2011). This decline in process rates can lead to a decrease in nutrient availability (Bengtson et al., 2005; Augusto et al., 2017). Conversely, the negative effects of water deficit on microbial processes may be partly compensated by increased activity following rainfall events (He and Dijkstra, 2014; Maxwell et al., 2022a). While drier soils can present slower microbial activity, the effect of increasing soil moisture may be constrained by soil nutrient availability (Cusack et al., 2019). In addition to affecting microbial processes, water availability can influence plant nutrient uptake and ecosystem nutrient retention. For example, lower precipitation may exacerbate the effect of N inputs, leading to greater plant N uptake, soil N availability and forest ecosystem N losses (Rivero-Villar et al., 2021). However, little is known about how limiting water conditions influence the expected beneficial effect of mixing tree species on nutrient availability, uptake, and cycling processes in forest ecosystems.

In this study, our main objective was to test the interactive effect of tree species mixing and water availability on a series of belowground nutrient cycling processes, in order to elucidate potential seasonal complementarity at varying temporal resolutions. To do so, we took advantage of the ORPHEE experimental site in southwestern France, a unique experimental design that manipulates *in situ* both the mixing of functionally different tree species (in this study, birch and pine) and water availability (*via* irrigation). We hypothesized (H_1_) that irrigation would lead to an overall positive effect on process rates, due to its favoring of microbial activity (Brockett et al., 2012). We expected this effect to be strongest during the dry summer period, when differences in soil moisture between irrigated and control plots would be greatest. We further hypothesized (H_2_) that species asynchrony between species would promote nutrient cycling, leading to increased organic matter mineralization (Borken and Matzner, 2009) and higher soil nutrient availability in mixed stands compared to monocultures (Zhang et al., 2026). More specifically, we expected mixing effects on soil nutrient availability to be strongest during the growing period, when microbial activity is highest (Davidson et al., 2006). Finally, we hypothesized (H_3_) that differences in phenology between birch and pine would generate temporal asynchrony in nutrient uptake, with the strongest differences occurring at budburst (Augusto et al., 2014). Because pine maintains functional foliage year-round whereas birch must first develop a new canopy, we expected nutrient uptake to be greatest at budburst and to differ between monocultures and mixtures.

## II Materials and methods

### Site description

The ORPHEE experiment is located in southwestern France (44°44.35’ N, 00°47.9’ W, elevation of 60 m asl.; Castagneyrol et al., 2013). The mean annual temperature is 13.2°C and the mean average precipitation is 940 mm (2000-2025), but most precipitation occurs from the fall to the spring (on average, 642 mm from October to April), often leading to dry summers considered representative of low water availability in this region. Plantation occurred in 2008, after a clear-cut of a *Pinus pinaster* forest with a single plough and fertilizer application of 26.2 kg of P and 49.8 kg of K per ha. The predominant soil type is a podzol characterized by a coarse texture (95% sand) and low soil fertility (Augusto et al., 2010), particularly in phosphorus (Trichet et al., 2009). The very coarse texture of the soil negatively affects its water holding capacity with values that is typically only ∼50 mm down to 1 meter (Augusto et al., 2006; Jolivet et al., 2007), with soil exploration by roots that is often limited by the presence of a hardpan (Achat et al., 2008) and anoxic winter conditions (Deirmendjian et al., 2018; Augusto et al., 2025). A total of 256 plots were planted, with five species (*Betula pendula* Roth., *Pinus pinaster* Soland. in Aït., *Quercus ilex* L., *Quercus pyrenaica* Willd., *Quercus robur* L.) in all possible combinations from one (monocultures) to five species mixtures. We chose to work with three species composition treatments: pure silver birch (*Betula pendula*) plots, pure maritime pine (*Pinus pinaster*) plots, and two-species mixed birch-pine plots (used to study the effect of tree mixing). Pine and birch were selected because, contrary to the other tree species, they grew more successfully under the harsh conditions imposed by understory competition, dry summers, and the low fertility of the sandy soil, resulting in relatively dense stands (Morin et al., 2020). Each plot has 100 trees within a 20 m by 20 m area. Trees were arranged in a substitutive (same tree density irrespective of the composition) alternate (trees from one species have neighbors from all other associated species) pattern. The plots are separated by 3 m from one another, and randomly located within six blocks: three control (dry) blocks and three irrigated blocks. The irrigation was started in 2015, and occurs seasonally from early May to late October/early November to alleviate summer water stress. The water is taken directly from the groundwater, which has low concentrations of nitrogen and phosphorus, but higher concentrations of other elements, such as calcium and magnesium (Augusto et al., 2025). A two-meter-tall sprinkler installed in the center of each experimental plot sprays 3 mm per night. On average, control blocks received 318 mm of water from early May to late September, while irrigated blocks received 777 mm of water (rain + irrigation) during the same period. When using both temperature and precipitation (+ irrigation) to determine arid periods in an ombrothermic diagram (Walter and Lieth, 1967), control blocks are subjected to more stress than irrigated blocks (Figure S1).

### Stand biomass measurements

Aboveground biomass of trees, and the understory, were estimated in 2018. The tree diameter at 1.3 m aboveground for all living trees within the central core area of the plot (i.e., excluding two perimeter ridges of trees), and allometric equations for pine (Shaiek et al., 2011) and birch (Johansson, 1999), were used to calculate an average biomass for each species in their monoculture and mixture. The understory biomass was measured by a phytovolume method (Gonzalez et al., 2013; Vidal et al., 2021). Briefly, each plot was divided into four zones, and the following present understory species were quantified: *Pteridium aquilinum L. Kuhn*, *Molinia caerulea L. (Moench)*, *Calluna vulgaris L. Hull*, *Erica cinerea L.*, *Erica scoparia L.*, *Frangula alnus P. Mill.*, *Ulex europaeus L.*, *Ulex minor Roth.*, *Rubus fruticosus L.*, and *Lonicera periclymenum L.* Their aboveground cover percentage was estimated, and average height and maximum height were measured (Vidal et al., 2021). We then calculated a sum of all understory aboveground biomass for each zone, and calculated an average per plot from the four zones.

### Experimental overview

The study of the nutrient availability and uptake was based on a three-approaches design. We placed (1) ion-exchange resins (IER), which captured the long-term nutrient accumulation over four periods of 6 months (hereafter referred to as “growing season” and “dormant season”, with May and November as start/end months), from November 2017 to November 2019. In order to better understand the mechanisms behind nutrient accumulation in the resins during the vegetation period, we also installed (2) *in situ* soil incubations following Raison’s method (Raison et al., 1987) to estimate net mineralization rates during shorter incubation periods from April to September 2019. Finally, we applied (3) a ^15^N labeled solution to the soil in order to estimate nutrient uptake at budburst 2020 and at peak phenology in early summer 2020. All of the methods are described below.

### Ion-exchange resins

Ion exchange resins (PST-1 Capsule, UNIBEST), were installed to estimate accumulated NH_4_^+^, NO_3_^−^ and PO_4_^3−^ in soils (Figure 1A). In each plot, four IER were placed in the top soil (at 8 cm deep, the 0 cm level being the limit between the top of the mineral layers and the bottom of the forest floor) on November 11^th^ 2017, and left for six months in the field with a metal grid to prevent wild boars from disturbing the soil, enabling the accumulation of available nutrients with a sensitivity to on-site factors (Binkley and Matson, 1983). The extraction of the resins allowed for the quantification of accumulated available N and P during two periods of the year, in the dormant season (≈ Oct./Nov. – April/May period) and the growing season (≈ April/May – Oct./Nov. period). On April 3^rd^ 2018, they were collected and new resins placed nearby. The IER were extracted by 1 M KCl and analyzed by colorimetric determination (SKALAR, San ++). This was repeated three times (analysis and installment of new resins on November 21^st^ 2018, and May 15^th^ 2019, finishing on October 16^th^ 2019). Results were standardized to correspond to µg capsule^−1^ 6 months^−1^ for each incubation period. Mineral N is hereto referred to as the sum of N-NH_4_^+^ and N-NO_3_^−^.

**Figure 1.**
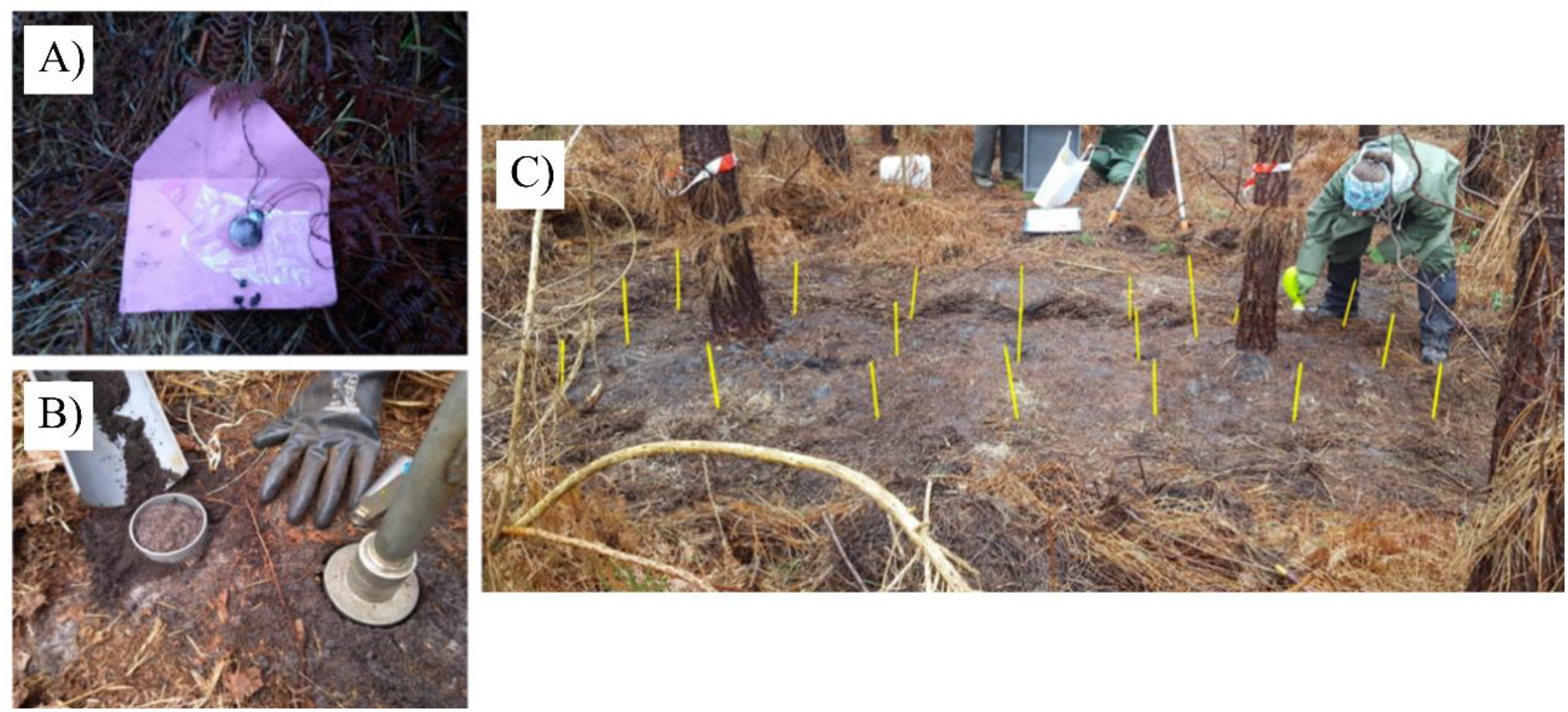
Images of the three methods and the experimental site. A) An ion-exchange resin after removal, B) incubation tube and soil corer for non-incubated soil analyses, and C) soil labeling using a small watering can in measured quadrants within a 3 m² zone (photo of T. Maxwell).

### Raison’s in situ soil incubation method

To study soil process rates, we installed *in situ* incubation tubes to estimate net mineralization rates in all of the studied plots (Raison et al., 1987, Figure 1B). This well-established method is able to detect small short-term (monthly to seasonal) change and cumulative effects of varying managements on soil mineral N dynamics (Khanna and Raison, 2013). The monitoring was carried out in spring-summer 2019, which is a period representative enough to capture the whole year dynamics (Schaffers, 2000).

Briefly, a set of undisturbed soil in tubes (depth: 15 cm, diameter: 8 cm) was installed in the soil and kept covered with a lid during field incubation. The mineral nutrient contents were measured in the unconfined soil at the time of installation of the incubation tubes. After around 5–8 weeks (exact dates detailed below), the mineral nutrient contents in both covered tubes and the unconfined soil was measured by water extraction and colorimetric analysis (SKALAR, San ++). The net rates for each nutrient [Nut] were calculated as below, with T2 corresponding to soil after the incubation and T1 to soils measured prior to incubation.

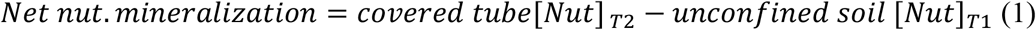

At the harvest date, the tubes were reinstalled nearby but in an undisturbed area, and the net mineralization was calculated again after another 5–8 weeks period (incubation times differed, but rates were standardized to 30 days^−1^ in order to be comparable between each other). This was repeated for a total of four times, in order to have estimations of net mineralization rates at four periods from early spring to late summer, corresponding to different soil conditions (Figure S2): fresh and moist soils with moderate microbial activity (April 1^st^ – May 9^th^), warm and still humid soils with strong activity (May 9^th^ – June 17^th^), dry soils with low activity (June 17^th^ – August 21^st^), and even drier soils during the fourth period (August 21^st^ – September 26^th^). Three field replicates were set up in each plot, from which the soil was homogenized in the field to produce one sample type per plot at each date. Samples were stored at 4 °C until analysis in the subsequent days.

Fresh soils were sieved to 2 mm, homogenized, and 1 aliquot per tube type per plot was extracted at each date (3 tube types × 3 species compositions × 2 water treatments × 3 blocks per water treatment = 54 samples per date). Briefly, 50 mL of ultrapure water was added to a glass vial with the 10 g aliquot, which was then shaken for 16 h at 4 °C. Water extraction was favored over KCl extraction due to potential interference of the extractant with available phosphate in sandy podzols (Achat, 2009), and because it has a similar extraction efficiency for mineral N as in saline solutions (Gallet-Budynek, INRAE; personal communication). The nutrients (NH_4_^+^, NO_3_^−^ and PO_4_^3−^) were analyzed in the extractant by colorimetric determination (SKALAR, San ++). Rates were calculated for each nutrient, and then the sum of the N-NH_4_^+^ and N-NO_3_^−^ were summed to produce one overall ‘net mineral N’ value.

### ^15^N soil labeling

Two pairs of trees were selected per plot, one pair for the budburst labeling and the other for the summer labeling. The surrounding aboveground understory was cleared in an area of 3.5 m × 1.5 m around the pair of trees, which were 2 m away from one another. After the soil was bare, a wooden quadrant and plastic sticks were used to accurately measure the area to be labeled (3 m × 1 m) around each pair of trees per plot (Figure 1C). The labeling solution was prepared with 55 mg of ^15^NH_4_Cl in 20 mL of deionized water in the laboratory. At the field, this was further diluted to 2 L of deionized water. The solution was sprayed over the labeling surface (3 L of 0.075 M, a non-fertilizing dose, over 3 m²), so that the soil was labeled with 7.5 mg ^15^N per m². The 3 L were sprayed evenly, by spraying 250 mL at a time, in each of the 12 measured squares of 50 cm × 50 cm using a small watering can (Figure 1C). Labeling occurred on March 16^th^ 2020 around budburst, and on June 8^th^ 2020, on the second pair of trees per plot, located at least 5 m away from the first pair labeled in March.

In August 2019, green leaves and new needles from several inner trees were sampled in order to get a base-line δ^15^N value for each species in each plot prior to the labeling experiment happening right around budburst (Figure S3). In March 2020, new leaves/buds and needles for each of the two trees in the labeling area were collected at two locations in the upper canopy using an extendable pruning shear, 1 week and 3 weeks after the labeling. For the second labeling date, green leaves were sampled in June 2020 a few days before the labeling occurred, to account for potential contamination from the previous nearby labeling experiment. The green leaves were also sampled 1 and 3 weeks after labeling. At all sampling dates in 2020, both the new needles and the 2^nd^ cohort (which had grown in 2019) were sampled. The new cohort from 2020 are included in the analyses and presented in this paper, and the values from the 2^nd^ cohort are presented in Figure S4. Leaves and needles for each tree at each sampling date were placed in individual paper bags and dried in an oven at 40 °C for at least 72 hrs. One composite was made from the two trees of each monoculture, so that per block there was 1 birch sample from its monoculture, 1 pine sample from its monoculture, 1 birch sample from the mixture, and 1 pine sample from the mixture. Each composite was cut up and ground, until the sample became a homogeneous powder. All material was wiped down with ethanol between samples to avoid contamination. A 2 mg aliquot of the powder was then weighed into tin caps on a microgram balance. The samples were analyzed for their δ^15^N using an Elemental Analyzer – Isotopic Ratio Mass Spectrometer (EA-IRMS), in the SILVATECH platform (Structural and functional analysis of tree and wood Facility) from the research center INRAE Grand-Est Nancy.

### Statistical analyses

All of the statistical analyses were done using R software (version 4.1.0). First, extreme values were removed following the outlier labeling rule with a conservative tuning parameter of g = 2.2 (Hoaglin and Iglewicz, 1987). On average, less than 5% of the IER values were removed, and no values were removed from the aboveground biomass, Raison’s method, nor the δ^15^N uptake data.

Mixed models were used to assess the effect of the following fixed effect factors: tree species composition (pure birch, pure pine, mixed birch-pine), water availability (irrigation, control), and their interaction, on the available nutrients extracted from the IER, mineralization rates, and understory biomass. Pairs of control and irrigated blocks (i.e., one ‘Superblock’) were included as a random effect to account for the spatial structure of our experimental design, with water availability treatment and Plot nested into nested into ‘Superblock’ (for IER only, as there were replicates per plot). For the ion-exchange resins and the mineralization rates, the *corAR1* autocorrelation function was used to take into account the correlation between the four time points, with date as the random slope and the spatial random effects as the random intercepts. The δ ^15^N dataset was separated into the two labeling dates, as individuals differed between the budburst and summer labeling experiments, and between pine and birch. A mixed-effect model was used to test the effect of mixing (monoculture, mixture), irrigation (irrigation, control), and their interaction, as well as of the sampling date (pre-label – only included for the summer experiment; post 1-week, and post 3-weeks). For the aboveground biomass data, a mixed-effect model was used to test the effect of mixing, species, irrigation, and their interactions. Both of these models included a spatial random effect (Plot nested in water availability treatment, nested in Superblock).

All mixed-effect models were coded using the ‘*nlme*’ package (Pinheiro and Bates, 2020) with α = 0.05 as significance level. Variables were log-transformed or arcsine transformed if needed to improve the normality of residuals. For all three data types and model results, we then applied contrasts and performed Tukey’s HSD tests using the ‘*multcomp*’ package (Hothorn et al., 2008), with α = 0.05 used as the level to define significance. Data was organized using the ‘*dplyr*’ package (Wickham et al., 2021) and graphs were produced with the ‘*ggplot2*’ package (Wickham, 2016).

## III Results

### Aboveground biomass

In general, pine had significantly higher tree aboveground biomass than birch (p < 0.001, Table 1, Table S1). Mixing birch and pine led to an overall positive non-additive effect on biomass in both control and irrigated plots (p < 0.001, Table 1, Table S1). While the theoretical additive value of mixing in control plots was 5.84 ± 1.26 kg m^−2^ (mean ± s.e.), the observed tree aboveground biomass was significantly higher, at 8.23 ± 0.75 kg m^−2^ (Table 1). Under irrigation, the difference was smaller from the theoretical additive effect (7.22 ± 1.2 kg m^−2^) to the observed effect (8.81 ± 0.11 kg m^−2^, Table 1). Irrigation had an overall positive effect on tree aboveground biomass (p = 0.004, Table S1).

**Table 1.**
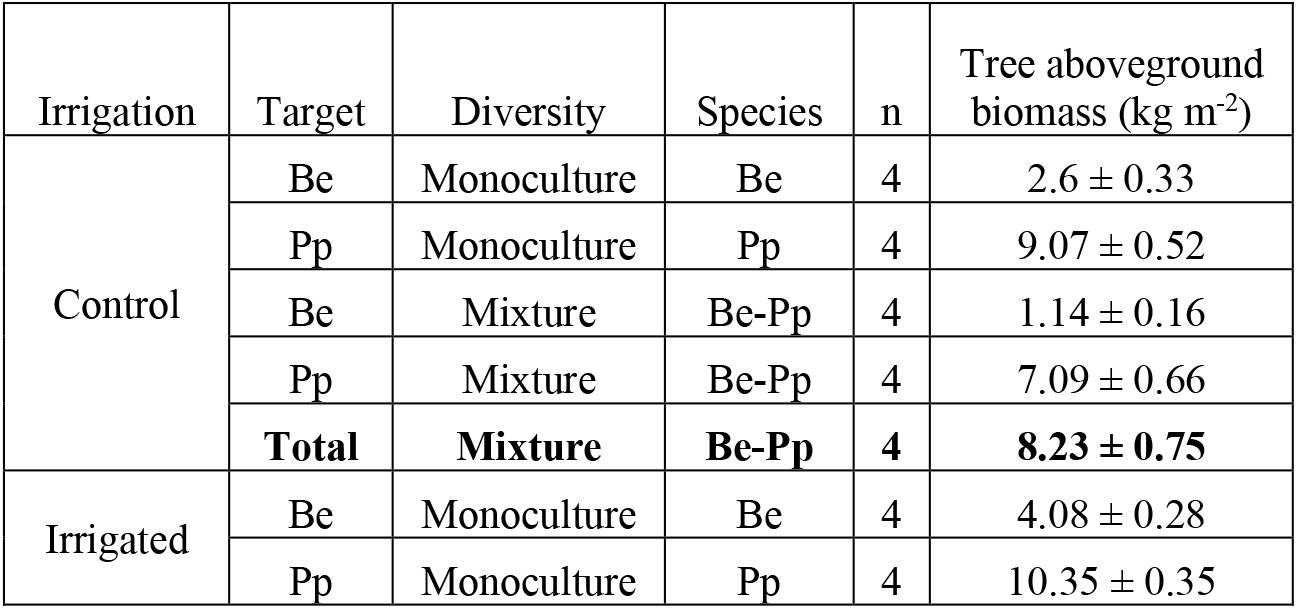
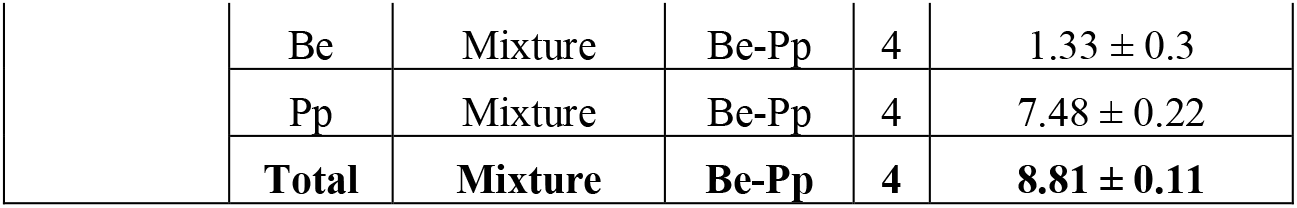
Tree aboveground biomass for birch (Betula pendula, Be) and pine (Pinus pinaster, Pp) grown in their monoculture and in the two-species mixture, under both control and irrigation water availability treatments. Values are mean biomass per tree per m² (± standard error), with measures taken in 2018. For each irrigation treatment, the total biomass per tree per m² (sum of birch and pine biomass) is presented in bold font.

The understory aboveground biomass at our experimental site significantly differed between species composition treatments (p< 0.001, Table 2, Table S1). Pure birch had three times more understory biomass than pure pine, in both control and irrigated plots (Table 2). The understory accounted for around 2% of total aboveground biomass (sum of understory and tree aboveground biomass) in pure birch plots, less than 0.25 % in pure pine, and 0.35 % in the mixed birch-pine plot.

**Table 2.** Understory aboveground biomass in each of the treatments (mean ± standard error), estimated in kg per m² for measures taken in 2018.

| Irrigation | Diversity | Species | n | Understory aboveground biomass (kg m<sup>-2</sup>) |
|---|---|---|---|---|
| Control | Monoculture | Be | 4 | 0.068 ± 0.010 |
| Control | Monoculture | Pp | 4 | 0.026 ± 0.009 |
| Control | Mixture | Be-Pp | 4 | 0.030 ± 0.004 |
| Irrigated | Monoculture | Be | 4 | 0.068 ± 0.007 |
| Irrigated | Monoculture | Pp | 4 | 0.023 ± 0.006 |
| Irrigated | Mixture | Be-Pp | 4 | 0.027 ± 0.004 |

### Soil nutrient availability

Overall, mineral N extracted from the ion exchange resins (IER) was affected by both species composition (p = 0.039) and by irrigation (p < 0.001, Table S1, Figure 2A). In general, pine tend to have higher mineral N captured by the IER, on average with 59 µg compared to 57 µg in the mixed-species plot and 33 µg in the pure birch plots (all units in µg capsule^−1^ 6 months^−1^). When considering all species compositions together, N in irrigated plots was reduced by 63 % compared to control plots (Figure 2A).

**Figure 2.**
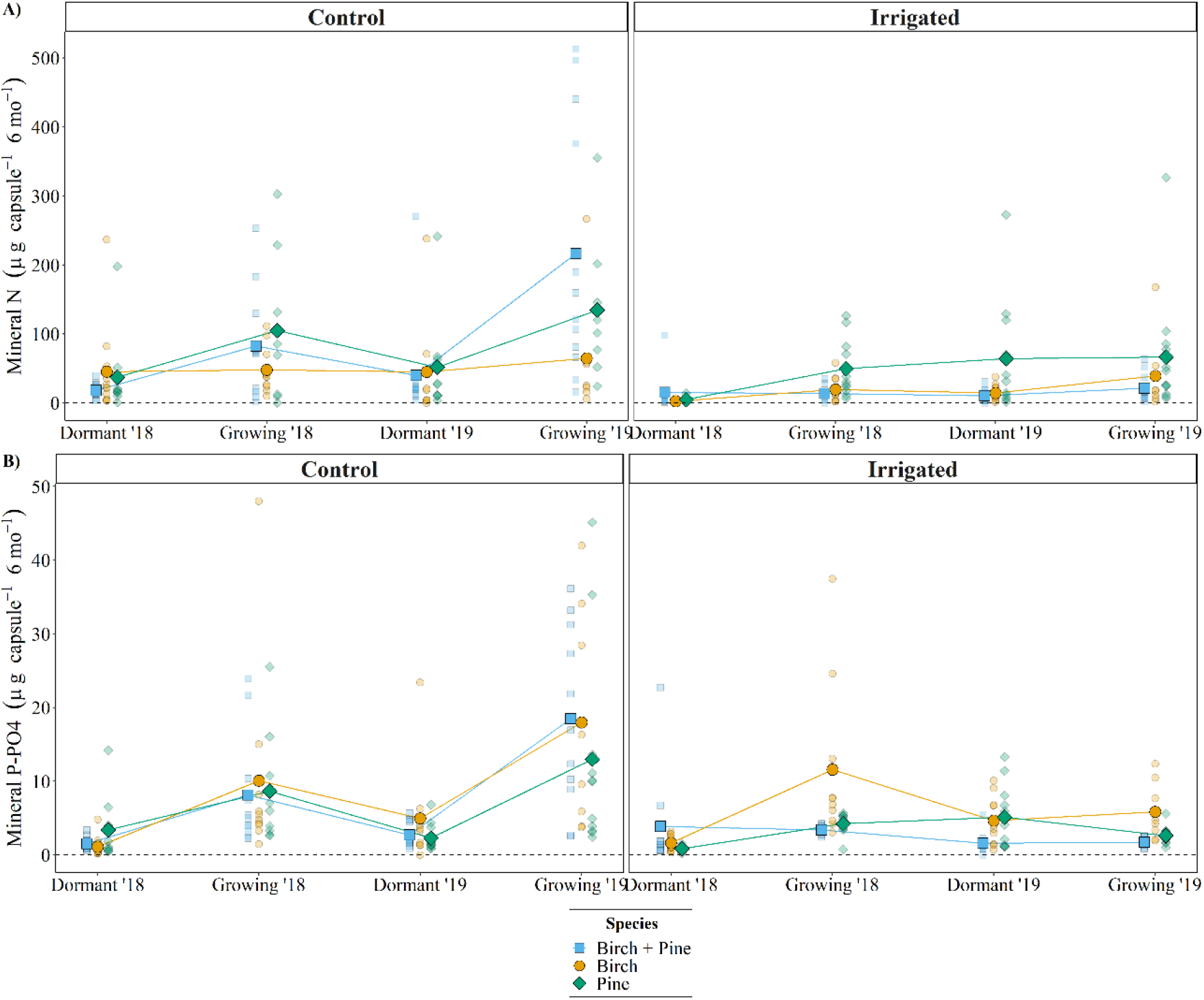
Soil mineral N (N-NH ^+^ + N-NO ^−^) and P-PO ^3−^ concentrations in mixed birch + pine (squares, *blue*), pure birch (circles, *yellow*), and pure pine plots (diamonds, *green*) under both control and irrigated water availability treatments. At each incubation period (dormant and growing seasons of 2018 and 2019), the raw data are presented in partial translucence, with the mean value superposed (n = 12).

Mineral P extracted from IER was negatively affected by irrigation (p = 0.011), on average by 46 % (Table S1, Figure 2B). There was also a significant interactive effect between irrigation and species on mineral P (p = 0.036). When sub-setting for each water availability treatment, birch had significantly higher P than both pine (p = 0.024) and the birch-pine mixture (p = 0.001) under irrigation (Figure 2B). Overall, this corresponded to 84 % more P available under birch than in pure pine, and 123 % more than in the mixture in the irrigated plots. Under control plots, no significant differences between species composition treatments were observed. For both N and P, models including season as a fixed factor revealed a significant effect of this effect (p < 0.001), with more nutrients recovered during the growing season than during the dormant season (Figure 2).

### Net nutrient mineralization

No significant effects of species composition, or its interaction with water availability, were found when analyzing the net nutrient mineralization rates estimated with Raison’s *in situ* incubation method (Table S1, Figure 3). Net nutrient mineralization values above 0 indicated net mineralization and negative values indicated net immobilization by soil microorganisms. Overall, there was a slight negative effect of irrigation on net N and P mineralization, i.e. nutrients were generally more immobilized than mineralized in irrigated compared to control plots (p = 0.016 for N and p = 0.013 for P, Figure 3). The patterns between the net mineralization of N and P were almost identical, with net N mineralization/immobilization being on average around three times greater than net P mineralization/immobilization (Figure 3). On average, N mineralization rates ranged from 0.64 ± 0.24 mg N kg^−1^ month^−1^ (mean ± s.e.) in control plots to –0.26 ± 0.23 mg N kg^−1^ month^−1^ in irrigated plots (Figure 3A). Phosphorus mineralization rates ranged from 0.20 ± 0.07 mg P kg^−1^ month^−1^ in control plots to –0.09 ± 0.07 mg P kg^−1^ month^−1^ in irrigated plots (Figure 3B).

**Figure 3.**
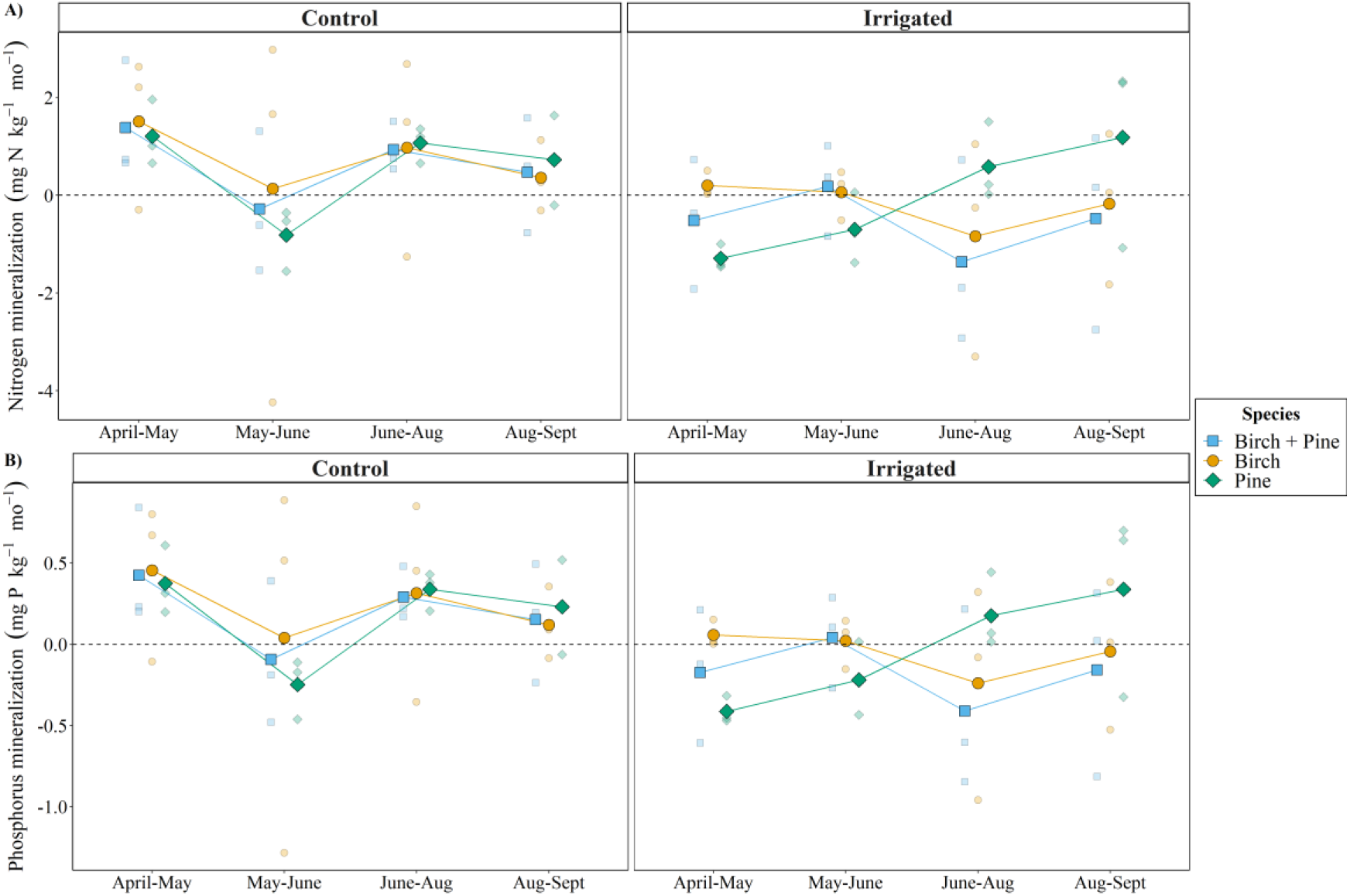
Mineral N and P mineralization in mixed birch + pine (squares, *blue*), pure birch (circles, *yellow*), and pure pine plots (diamonds, *green*) under both control and irrigated water availability treatments. Nutrient mineralization values above 0 indicate net mineralization, and values below 0 indicate immobilization. At each incubation period in 2019, the raw data are presented in partial translucence, with the mean value superposed (n = 3).

### Uptake

Throughout the labeling experiment, pine needles generally had higher δ^15^N values than birch leaves (Figure 4). This was also true for the leaves sampled in summer 2019, prior to the budburst labeling experiment, for which birch had a δ^15^N value of –0.87 ± 0.28 (mean ± s.e.) and pine had higher values, around 6.82 ± 0.65 δ^15^N (Figure S3). There were no differences between treatments for each species (Figure S3).

**Figure 4.**
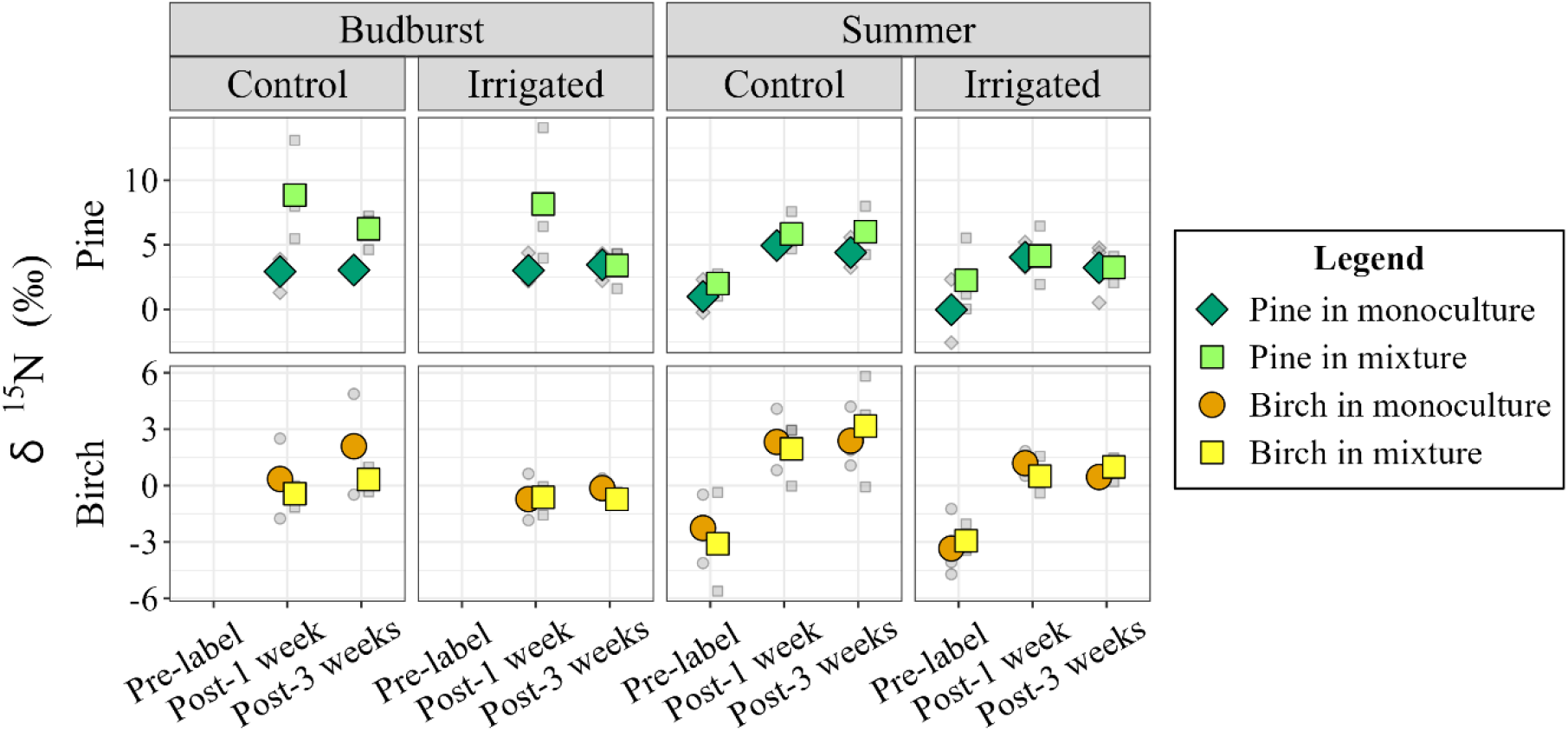
Values of pine needle (top) and birch leaf (bottom) δ^15^N prior to labeling, 1 week after labeling, and 3 weeks after labeling, for the budburst (March 16^th^) and summer (June 8^th^) 2020 soil labeling campaigns. Pre-labeling sampling for the budburst campaign occurred in July 2019 and was not included in this analysis (Fig. S3). Data are presented for pine in its monoculture (diamonds, *green*) and in the birch-pine mixture (squares, *light green*), and for birch in its monoculture (circles, *yellow*) and in the birch-pine mixture (squares, *light yellow*). At each leaf sampling date, the raw data are presented in *grey* and partial translucence, with the mean value superposed (n = 3).

After the budburst labeling, the new pine cohort in its mixture had significantly more δ^15^N than in its monoculture (p = 0.018, Table S1), with values one week after labeling around 8.51 ± 1.70 δ^15^N in the mixture and 2.98 ± 0.48 δ^15^N in the monoculture (Figure 4). This effect was not observed for birch, with values that centered around 0 δ^15^N (Figure 4). The second cohort of pine needles also did not show differences between treatments (Figure S4). Values were similar three weeks after labeling, suggesting that the label had mainly been taken up in the first week.

During the summer labeling experiment, the effects of mixing were not observed; both species behaved similarly in their monoculture and in the mixture (Figure 4). Both species also showed a strong increase in δ^15^N one-week after labeling, a signal that stayed constant at the post-three-week sampling. On average, pine δ^15^N increased by 264 % one week after labeling, while birch increased by 151 % compared to the pre-labeling sampling date a few days prior. There was no overall effect of irrigation on the 15N uptake for either species at either labeling date (Table S1).

## IV Discussion

### Effect of water availability on belowground processes

Contrary to our first hypothesis (H_1_), belowground process rates did not accelerate with an increase in water availability. Over the growing period, irrigation tended to have a negative effect on net nutrient mineralization, indicating higher net microbial immobilization in these plots (Zhang et al., 2018). This could be due to the higher competition between microbes and trees (Laungani and Knops, 2012), as irrigation increased aboveground productivity (Table 1). As for tree nutrient uptake, the δ^15^N signal was not stronger in irrigated blocks as there was no difference between water availability treatments. Although this is consistent with the late activation of the irrigation treatment in 2020 (one week prior to labeling), the lack of water treatment effect implies that there was not a strong legacy effect of reduced water availability on nutrient uptake. Contrary to previous studies finding an increased allocation to roots with drier conditions (Bakker et al., 2009; Bello et al., 2019), our site did not exhibit differences in root biomass between control and irrigated blocks (Altinalmazis-Kondylis et al., 2020); this may explain why we observed no difference in uptake with water availability. Additionally, there was a negative effect of irrigation on available mineral N and P, the latter being most affected by irrigation during the growing season (i.e. when irrigation was in effect; Figure 2). Leaching resulting from increased water inputs may explain this overall negative effect. The ion-exchange resins were placed 8 cm belowground, and a previous study at our site found that irrigation decreased available nutrients in the top 0-15 cm layer, but then increased nutrients down to 90 cm (Maxwell et al., 2020b). Overall, our results indicate that increasing water availability modified nutrient cycling processes, but not in the direction initially expected. Rather than stimulating nutrient availability and uptake, irrigation appeared to promote nutrient immobilization and redistribution within the soil profile.

### No overall effect of mixing on soil nutrient availability

The soil incubations using Raison’s method to estimate net nutrient mineralization at various periods between early-April and late-September, showed no significant differences among species composition treatments (Figure 3). Notably, the trends observed for net N mineralization were the same as for net P mineralization, suggesting a stable microbial N:P stoichiometry in our soils (Mooshammer et al., 2012; Fanin et al., 2013). In general and as expected, more nutrients were recovered during the growing season, due to the stimulation of microbial activity with the temperature increase at this time (Davidson et al., 2006). These fluctuations between dormant and growing season were especially apparent in the control blocks (Figure 2), but this is likely due to the leaching in the irrigation treatment, as discussed previously. Over the various six-month ion-exchange resin (IER) incubation periods, there was no effect of mixing on soil nutrient availability, in line with the neutral effect of mixing on net nutrient mineralization (Figure 3). This may be because the IER captured nutrients that were neither immobilized by microorganisms, nor taken up by surrounding trees. In addition, interesting trends emerged among species composition treatments under improved water availability. Significantly higher mineral N was found in the pure pine than in either the pure birch or mixed birch-pine plots, and this was consistent over the two years of monitoring. The poor sandy soils in the experimental site’s region have a widespread and severe limitation in phosphorus (Trichet et al., 2008; Achat et al., 2009). Pine trees in the monocultures were around 2.5 times larger than the birch trees (Table 1). Because the growth rate in pure pine stands is high, the nutrient demand is high (Augusto et al., 2000, 2008), and thus the available P pool may be more limiting. This may alleviate the demand for N, which eventually enabled the N pool to increase in pure pine stands. Consistent with these findings, the pure birch plots under irrigation also displayed significantly higher available P than pure pine or the mixture during the growing season (Figure 2B). This is similar to a previous study on volcanic soils, which found that as precipitation increased, the effect of species was more relevant, with broad-leaved deciduous species exhibiting the highest values of P compared to evergreen conifers (Satti et al., 2007). Although each of the monocultures had different increases in nutrient availability, this was not combined or apparent in the mixture. We thus reject our second hypothesis (H_2_), as we did not observe an increase in net nutrient mineralization or soil nutrient availability with mixing. However, during the last growing season of the experiment, there was a trend for higher soil nutrient availability in the mixture. Therefore, future works with more time since plantation may reveal long-term nutrient complementarity in the mixture.

### Asynchrony of uptake at budburst, then lost at summer

In order to explore potential asynchrony of uptake between birch and pine in the mixture, soil labeling campaigns were done at budburst, and at peak leaf development in early summer. Prior to these campaigns, pine had a higher ^15^N isotopic signal than birch (Figure S3), although both species had values consistent within the range of values for the mean annual temperature and precipitation of our experimental site (Craine et al., 2015a) and for local vegetation (Augusto et al., 2005; Cavard et al., 2007). This difference may reflect contrasting sources of nutrient acquisition, as δ^15^N increases with depth (Craine et al., 2015b), a pattern which is also observed locally (Augusto et al., 2005). This suggests that pine may acquire a greater proportion of its nutrients from deeper soil layers, despite the similar root profiles observed for pine and birch at our site (Altinalmazis-Kondylis et al., 2020).

The labeling campaign at budburst revealed contrasting responses of the two species in the mixture. At this stage, pine displayed higher δ^15^N than in its monoculture, while birch’s δ^15^N value was similar to its monoculture. Although the isotopic signal does not allow us to quantify the total amount of N acquired by each species, it provides valuable information on their relative access to recently labeled soil N. At the current age and density, pine overtook birch and largely dominates the canopy and productivity of the mixture (Morin et al., 2020). After 12 years of growth, birch trees were overshadowed by pine (Table 1, Figure S5), which largely took up canopy space (Martin-Blangy et al., 2023). This canopy dominance has been associated with delayed budburst for birch in the mixture compared to its monoculture, as previously found for oak mixed with pine (Perot et al., 2021). While N absorption can occur before budburst (Bazot et al., 2016; Maxwell et al., 2020a), pine trees retains foliage from previous needle cohorts and may therefore begin transpiration and nutrient acquisition earlier than birch (Rothe, 1997). The stronger δ^15^N enrichment observed in pine growing in mixture suggests greater access to recently available soil N during this period, whereas birch appeared to benefit less from the labeled N pool. This pattern is consistent with a temporary release from intraspecific competition for pine combined with stronger competition experienced by birch in mixture, creating a transient win-lose situation (Forrester et al., 2011; Nanami et al., 2011; Vanclay et al., 2013). However, because δ^15^N enrichment reflects access to labeled N rather than total N acquisition, these results should be interpreted cautiously. Moreover, interpretation of pine δ^15^N dynamics requires consideration of needle cohort turnover. Needles sampled before and after the budburst labeling did not belong to the same cohort, and accounting for this cohort effect suggests greater continuity in δ^15^N values through time than would be inferred from direct comparisons among sampling dates (Figure S4). Pine enrichment relative to birch is unlikely to result from differences in understory biomass, as the latter was slightly higher in the mixture than in pure pine (Table 2). Additionally, the response was similar in both the irrigated and the control plots, which corroborates the lack of a legacy effect of differences in water availability on nutrient uptake. Overall, stand structure, including tree size, canopy occupation, and phenological differences between species, appears to have played a major role in determining nutrient uptake dynamics at budburst. The summer labeling campaign resulted in similar uptake patterns for the two species in their monocultures and mixture (Figure 4). By early June, both trees had reached their maximum leaf area, and likely similar nutrient capacities prior to onset of severe summer drought (Gholz et al., 1990). The contrasting enrichment patterns observed at budburst had therefore disappeared by early summer. These results support our third hypothesis (H_3_) that temporal asynchrony in nutrient uptake between birch and pine would be strongest at budburst.

### Synthesis and implications for nutrient cycling in mixed species forests

The combination of ion-exchange resins, *in situ* soil incubations and soil ^15^N labeling provided complementary insights into belowground nutrient cycling processes at different temporal scales (Figure 5). Contrary to our expectations, neither irrigation nor species mixing increased net nutrient mineralization, which was measured with *in situ* soil incubations. However, species composition influenced soil nutrient availability, with all pure pine stands exhibiting higher mineral N and the pure birch stands under high water availability exhibiting higher available P in the ion-exchange resins. The strongest evidence for temporal complementarity was observed during budburst. At this stage, pine exhibited greater δ ^15^N enrichment than in its monoculture following the ^15^N labeling experiment, while birch showed similar values to its monoculture. This asymmetry disappeared by early summer, when both species had reached their maximum foliage development and showed similar enrichment patterns following soil labelling between the mixture and their respective monocultures. These results suggest that phenological differences between species can temporarily influence access to recently available soil N. Taken together, our findings indicate that differences in nutrient availability among species compositions were not associated with differences in net nutrient mineralization. Instead, species-specific nutrient demand, nutrient uptake dynamics, and stand structure appear to play a greater role in determining nutrient availability.

**Figure 5.**
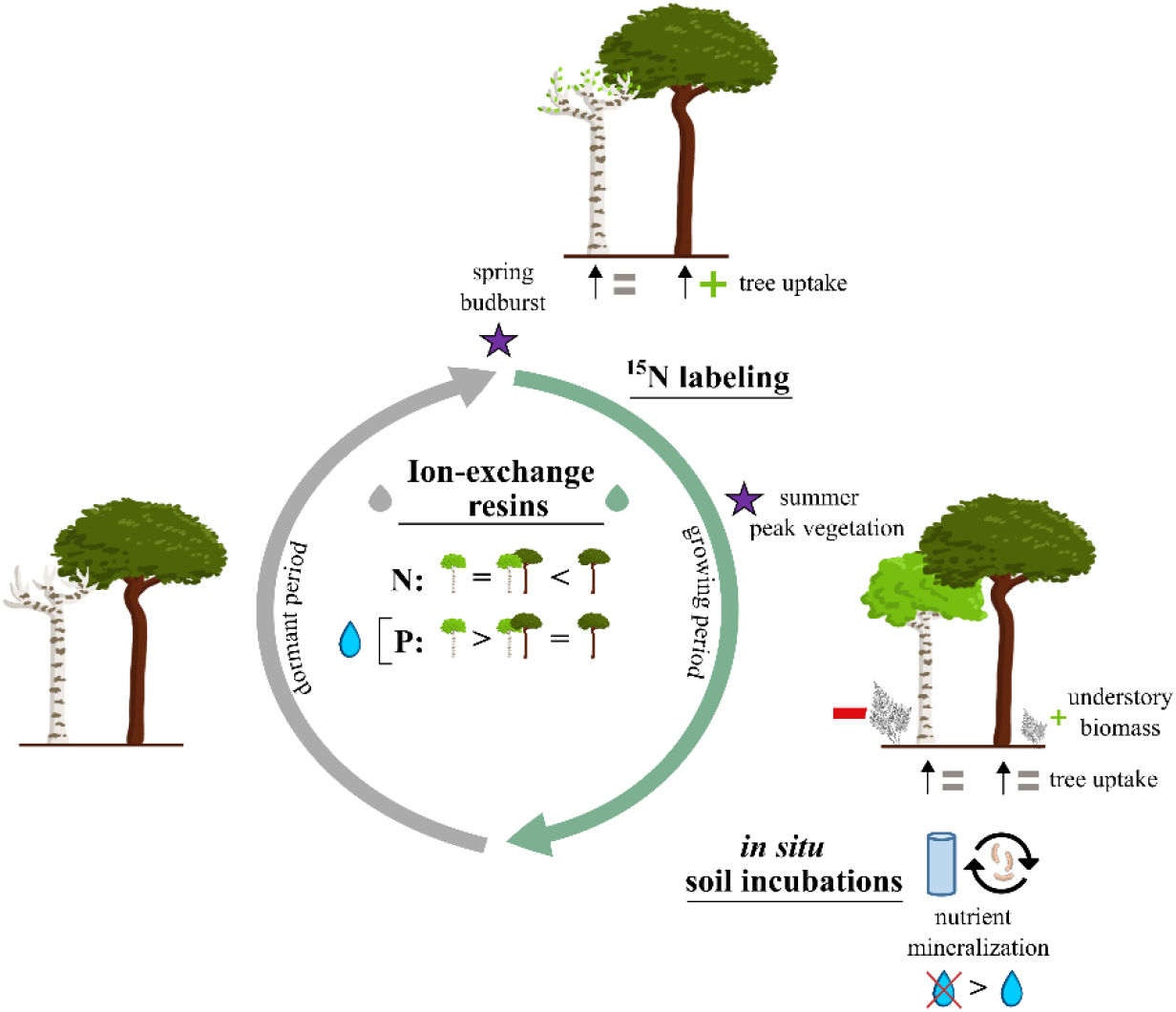
Belowground nutrient cycling at various periods throughout the year. (Top) From the spring budburst labeling campaign (*purple* star), birch took up similar nutrients to its monoculture while pine took up relatively more than its monoculture (*green* positive symbol). (Right) After the peak summer vegetation labeling campaign (*purple* star), both birch and pine in the mixture had similar uptake rates as in their respective monocultures. During summer, the mixture had greatly less understory than the pure birch plots (*red* negative symbol), slightly higher than pure pine (*green* positive symbol), and overall similar mineralization by soil microorganisms. Nutrient mineralization rates were overall higher in control than in irrigated plots. (Center) Under irrigation, more nitrogen (N) was available in pure pine than in pure birch and the mixture, while more phosphorus (P) was available in pure birch than either of the other two stands. The nutrient availability trends stayed constant from the growing to the dormant period.

## V Conclusions

Using a unique experimental design manipulating both tree species composition and water availability, we investigated the drivers of belowground nutrient cycling in mixed birch-pine stands. While there was no effect of species mixing nor irrigation on net nutrient mineralization, species composition influenced soil nutrient availability, especially under high water availability. This suggests that differences in nutrient demand among species play an important role in shaping belowground nutrient dynamics. Mixing tree species with different phenologies and growth rates influenced nutrient cycling through asynchrony in nutrient uptake at budburst and different levels of nutrient demand. More generally, our results suggest that temporal complementarity in mixed forests may arise primarily through differences in the timing of nutrient acquisition rather than through enhanced microbial mineralization, highlighting the importance of species phenology in regulating belowground nutrient cycling.

## Data and code availability

The data and code that support the findings of this study are openly available in Zenodo: https://doi.org/10.5281/zenodo.21640158

## Acknowledgements

First, we would like to thank Bastien Castagneyrol, Hervé Jactel, Bernard Issenhuth and the UEFP team for maintenance of the ORPHEE experiment. We also thank INRAE Nancy for δ^15^N analyses. We thank Laura Scolan for her help with the *in situ* incubations set up and analyses. We are grateful to Noé Augusto, Gwenola Augusto, and Lucie Bon for preparing the ion-exchange resins, and Aline Fanin for lending us her small watering can for the soil labeling experiment. Additionally, we thank Malika Aimaiti, Andreas Altinalmazis-Kondylis, Mark R. Bakker, Lucie Bon, Julien Hennequart, Marko Kvacic, Sylvie Niollet, and Pablo Raguet for help in the field. Finally, the help of Coralie Chesseron, Adèle Courbineau, Nathalie Gallegos, Sylvie Milin, and Pierre Pujos in the lab is greatly appreciated. This work was supported by the *Agence Nationale de la Recherche* (ANR), France project DiPTiCC (16-CE32-0003), and the “Thèses en partenariat international” project, Excellence Initiative, Université de Bordeaux, France to TLM. ADM supported TLM through an NSERC (Natural Sciences and Engineering Research Council) Discovery grant. We are deeply saddened by the burning of the ORPHEE experimental site during the July 2026 wildfires, and express gratitude the hard work of the firefighters and community members.

