## Supplementary Information for "Tree species composition and nutrient uptake influence soil nutrient availability, even under contrasted levels of water availability"

**Table S1** Results of the mixed model analyses of species composition (Sp), water availability treatment (Irr) and their interaction on understory aboveground biomass, soil nutrient availability and net mineralization rates. For the tree aboveground biomass and δ15N uptake values, results come from the mixed model analysis of diversity (Div), Sp, Irr, and their interactions. Numerator and denominator degrees of freedom, as well as F-values and p-values for fixed effects are presented. Significant ($p<0.05$) effects are noted in bold font.

| Variable | Fixed effect | numDF | denDF | F-value | p-value |
|---|---|---|---|---|---|
| Tree aboveground biomass | Div | 1 | 21 | 77.90 | < **0.001** |
| | Sp | 1 | 21 | 583.93 | < **0.001** |
| | Irr | 1 | 21 | 10.49 | **0.004** |
| | Div x Sp | 1 | 21 | 0.38 | 0.546 |
| | Div x Irr | 1 | 21 | 4.49 | **0.046** |
| | Sp x Irr | 1 | 21 | 0.00 | 0.994 |
| | Div x Sp x Irr | 1 | 21 | 0.16 | 0.696 |
| Understory aboveground biomass | Sp | 2 | 15 | 23.61 | < **0.001** |
| | Irr | 1 | 15 | 0.15 | 0.702 |
| | Sp x Irr | 2 | 15 | 0.03 | 0.973 |
| Mineral N ($N\text{-}NH_4^+ + N\text{-}NO_3^-$) | Season | 1 | 178 | 51.76 | < **0.001** |
| | Irr | 1 | 10 | 40.67 | < **0.001** |
| | Sp | 2 | 10 | 41.57 | **0.039** |
| | Irr x Sp | 2 | 10 | 1.23 | 0.333 |
| Mineral P ($P\text{-}PO_4^{3-}$) | Season | 1 | 184 | 116.89 | **<0.001** |
| | Irr | 2 | 10 | 9.49 | **0.011** |
| | Sp | 1 | 10 | 1.34 | 0.305 |
| | Irr x Sp | 2 | 10 | 4.71 | **0.036** |
| Net N mineralization | Sp | 2 | 10 | 0.22 | 0.803 |
| | Irr | 1 | 10 | 8.29 | **0.016** |
| | Sp x Irr | 2 | 10 | 0.29 | 0.754 |
| Net P mineralization | Sp | 2 | 10 | 0.28 | 0.762 |
| | Irr | 1 | 10 | 9.17 | **0.013** |
| | Sp x Irr | 2 | 10 | 0.27 | 0.768 |
| Budburst $^{15}N$ uptake Birch | Irr | 1 | 6 | 2.54 | 0.162 |
| | Div | 1 | 6 | 0.97 | 0.362 |
| | Date | 2 | 11 | 8.61 | **0.014** |
| | Irr x Div | 1 | 6 | 0.20 | 0.667 |
| Budburst $^{15}N$ uptake Pine | Irr | 1 | 6 | 0.62 | 0.461 |
| | Div | 1 | 6 | 10.41 | **0.018** |
| | Date | 2 | 11 | 1.74 | 0.214 |
| | Irr x Div | 1 | 6 | 1.43 | 0.276 |
| Summer $^{15}N$ uptake Birch | Irr | 1 | 6 | 2.99 | 0.134 |
| | Div | 1 | 6 | 0.00 | 0.972 |
| | Date | 2 | 22 | 84.60 | **<0.001** |
| | Irr x Div | 1 | 6 | 0.02 | 0.885 |
| Summer $^{15}N$ uptake Pine | Irr | 1 | 6 | 3.9 | 0.095 |
| | Div | 1 | 6 | 2.5 | 0.162 |
| | Date | 2 | 22 | 16.3 | **<0.001** |
| | Irr x Div | 1 | 6 | 0.1 | 0.771 |

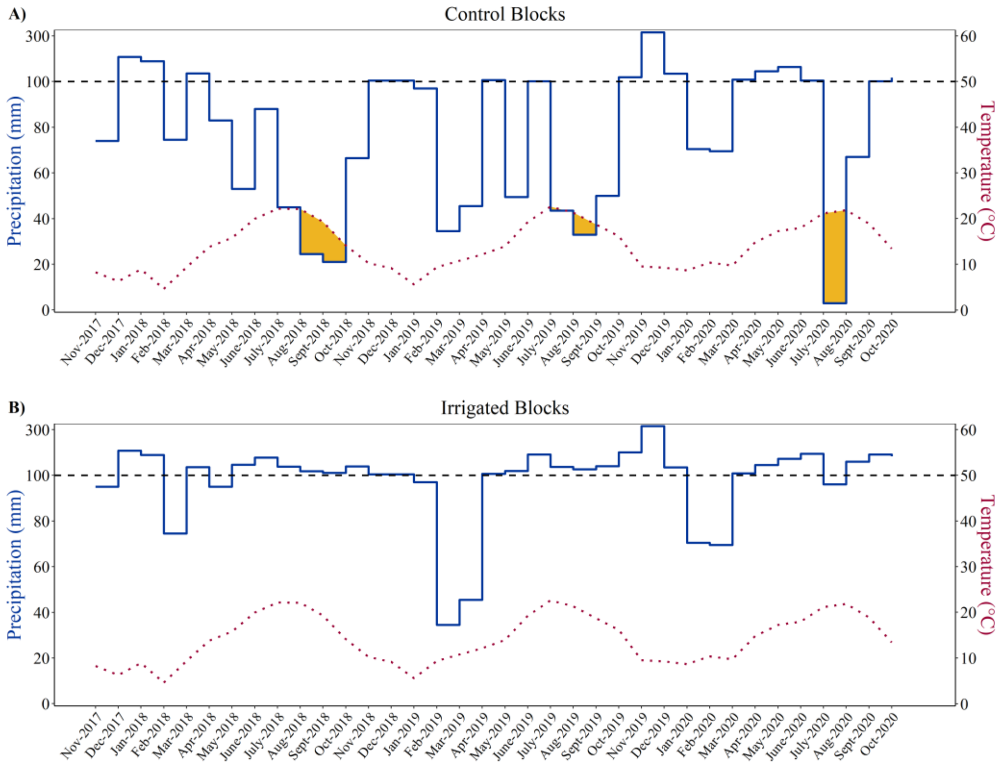


**Figure S1** Ombrothermic diagram with total monthly precipitations (blue solid line) and average monthly temperature (red dotted line) during the study period for the control and irrigated treatments. Data were obtained from an INRAE weather station at Cestas Pierroton (44.742 °N - 0.782 ° E), within 1 km of the experimental site. Temperature scale is set to 2mm/°C, and values above the 100mm black dashed horizontal line are scaled to 20mm/°C (Walter and Lieth, 1967). When temperature is two times greater than precipitation, there is an arid period identified in yellow.

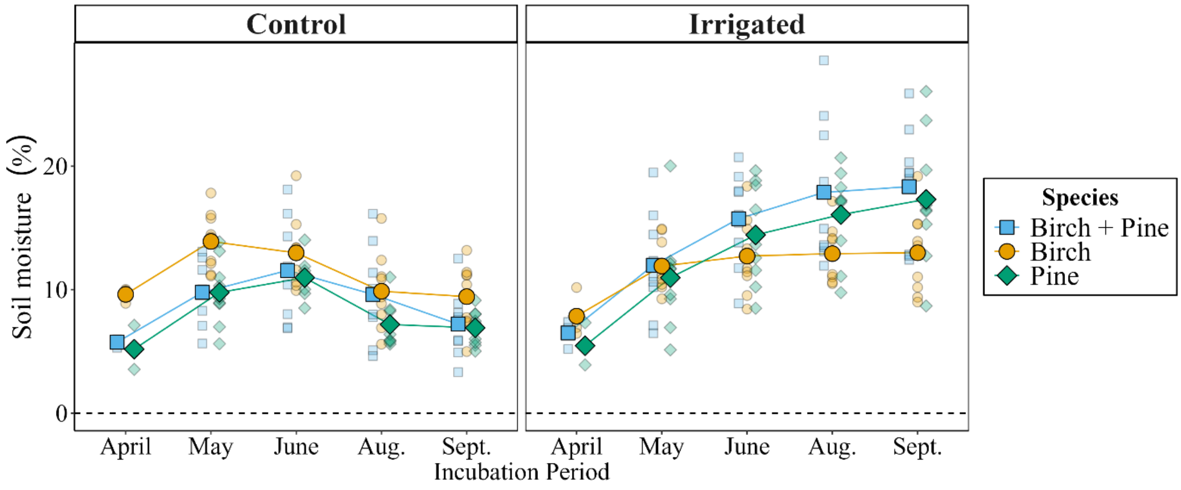


**Figure S2** Soil moisture from uncovered soils recovered at each sampling date during Raison's soil incubation method in 2019. Values for mixed birch + pine (squares, blue), pure birch (circles, yellow), and pure pine plots (diamonds, green) are presented under both control and irrigated water availability treatments. The raw data are presented in partial translucence, with the mean value superposed.

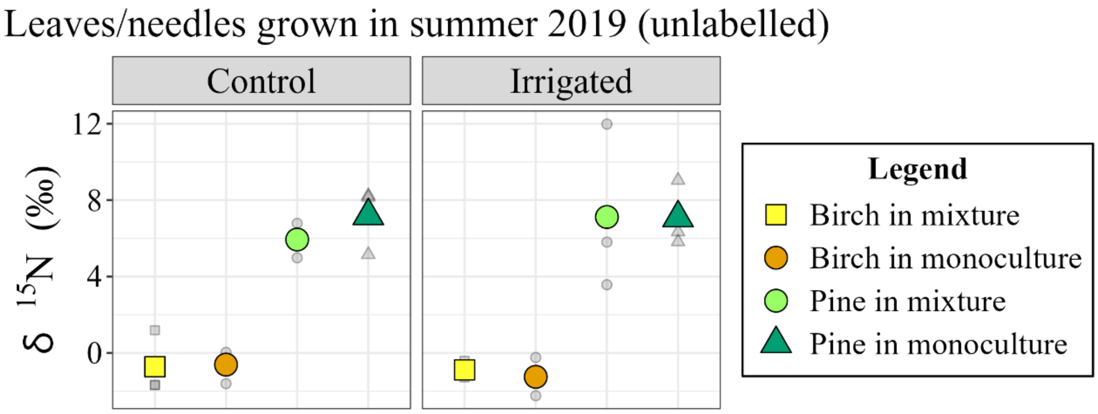


**Figure S3** The $\delta^{15}N$ values of birch leaves and new pine needles (1st cohort) sampled in August 2019, the summer prior to the $^{15}N$ labeling experiments. The raw data are presented in *grey* partial translucence, with the mean value superposed (n = 3).

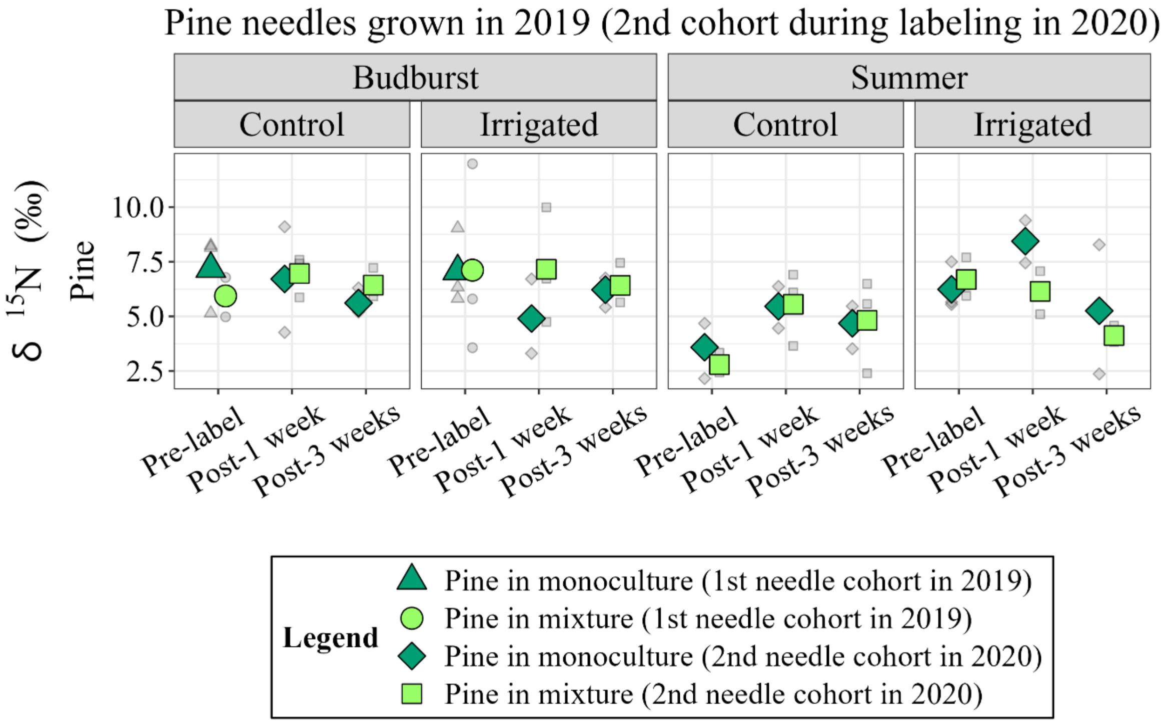


**Figure S4** The $\delta^{15}N$ values of new pine needles (1st cohort) sampled in August 2019 (pre-label for Budburst), which became the 2nd cohort during the labeling experiment in 2020 (Budburst: Post-1 week and Post-3 weeks; all dates in Summer). The raw data are presented in *grey* partial translucence, with the mean value superposed (n = 3).

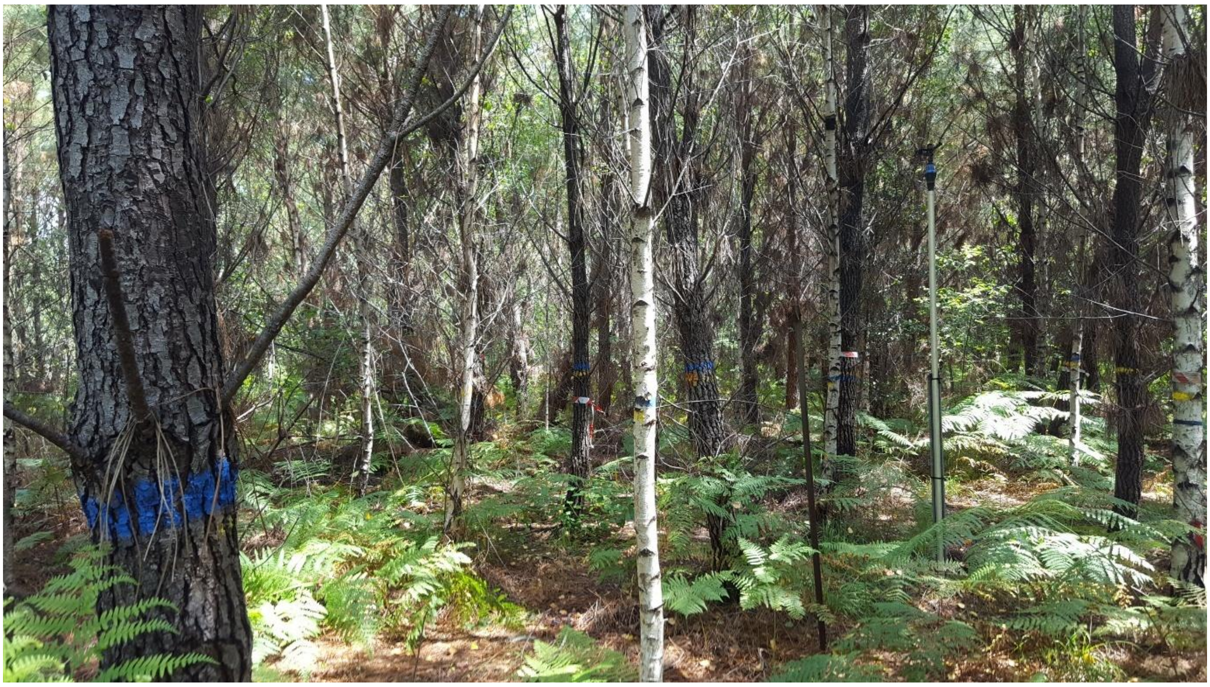

**Figure S5** A mixed birch-pine irrigated plot (photo taken in June 2020). Pine trees are larger than birch trees, offering a competitive advantage for nutrient uptake at budburst.
